# Refined Tamoxifen Administration in Mice by Encouraging Voluntary Consumption of Palatable Formulations

**DOI:** 10.1101/2023.04.24.538122

**Authors:** Dominique Vanhecke, Viola Bugada, Thorsten Buch

## Abstract

Drug administration in preclinical rodent models is essential for research and development of novel therapies. Compassionate administration methods have been developed, but these are mostly incompatible with water-insoluble drugs such as tamoxifen or do not allow for precise timing or dosing of the drugs. For more than two decades, tamoxifen has been administered by oral gavage or injection to CreER^T2^/loxP gene-modified mouse models to spatiotemporally control gene expression, with the numbers of such models steadily increasing in recent years. Animal-friendly procedures for accurately administering tamoxifen or other water-insoluble drugs would therefore have an important impact on animal welfare. Based on a previously published micropipette feeding protocol, we developed palatable formulations to encourage voluntary consumption of tamoxifen. We evaluated the acceptance of the new formulations by mice during training and treatment and assessed the efficacy of tamoxifen-mediated induction of CreER^T2^/loxP dependent reporter genes. Both sweetened milk and syrup-based formulations encouraged mice to consume tamoxifen voluntarily, but only sweetened milk formulations were statistically non-inferior to oral gavage in inducing CreER^T2^-mediated gene expression. Serum concentrations of tamoxifen metabolites, quantified using an in-house developed cell assay, confirmed the lower efficacies of syrup- as compared to sweetened milk-based formulations. We found dosing with a micropipette to be more accurate, with the added advantage that the method requires little training for the experimenter. The new palatable solutions encourage voluntary consumption of tamoxifen without loss of efficacy compared to oral gavage and thus represent a refined administration method.

## Introduction

The increased efforts to conduct more humane animal research (3R’s principle^1, 2^) include the development and application of animal-friendly drug administration methods. Unlike water-soluble drugs, most water-insoluble drugs cannot be mixed with drinking water or chow. Instead, they are typically administered via oral gavage (OG), injected intraperitoneally (IP) or subcutaneously (SC). These invasive interventions can induce stress-related responses in rodents, as reflected by increased stress hormone levels or heart rates^3–8^. Furthermore, repeated OG can increase the risk of unintentional injuries, including perforation of the trachea, esophagus or stomach, introduction of fluids into the trachea or lung, and hemorrhage^9^. Repeated IP injections have been reported to cause local irritation, pain, infection, and damage to surrounding tissue^10^.

Recently, a procedure for drug administration in mice that aims to minimize the above-mentioned disadvantages has been proposed as an alternative to OG or IP injections^7, 8^. This so-called micropipette-guided drug administration (MDA) makes use of a sweetened condensed milk solution as a vehicle to motivate mice to voluntarily consume drug solutions. This non-invasive procedure was shown to achieve pharmacokinetic profiles similar to those obtained by oral gavage ^7^ or IP injections ^8^. However, until now, similar pipette feeding has not been successfully adopted for the administration of water-insoluble compounds such as, for example, tamoxifen (TAM).

Tamoxifen is a selective estrogen receptor (ER) modulator that is widely used in clinical and basic research applications. For more than 20 years, tamoxifen has been utilized in research to induce spatiotemporal modifications in gene expression in Cre-ER^T^^1/T2^/loxP transgenic mouse models ^11–14^. The number of CreER^T1^ and CreER^T2^ mouse models generated for research exceeded 1,000 by June 2021^15^, and an additional 100 models were generated in the last year alone (as of May 2022)^15–17^. Despite its routine use in research, there is no consensus on the best method for TAM delivery^11^. In most cases, it is administered via OG or IP injections, but also occasionally by SC injections or medicated diets^18, 19^. While injections allow controlled dosing and timed treatments, the oil- or ethanol-based vehicles required to dissolve tamoxifen can cause local adverse reactions at injection sites ^10, 11, 20^. Oral administration is more physiologically relevant for tamoxifen since it first needs to be metabolized by the liver into the biologically active metabolites 4-hydroxytamoxifen and endoxifen^21, 22^. However, of the existing oral administration methods, OG is an invasive method that requires restraint of the animal and specific training by the experimenter. Treatment with TAM-supplemented diets on the other hand, while it is convenient, does not allow accurate dosing and can result in adverse effects from poor feeding due to aversion^23^.

Developing a more refined method for TAM administration would not only benefit the well-being of experimental mice, but it should also improve experimental outcomes. Indeed, prevention of possible stress-related confounder effects^24–27^ and increased accuracy of dosing and timing of treatments are expected to improve experimental conditions and thus the quality of the results. This, in turn, will reduce animal use overall, since smaller treatments groups are required to yield statistically significant results. A refined administration method could also benefit research involving other water-insoluble drugs such as Cyclosporin A or antibiotics, including Linezolid and Vancomycin, which are currently administered to mice or rats by OG or IP injections^28–30^. Developing palatable formulations for animal-friendly administration of water-insoluble drugs could even have a larger impact, considering that approximately 40% of drugs with market approval and nearly 90% of molecules in the discovery pipeline are poorly water-soluble^31^.

In this study we developed sweetened formulations that are compatible with water-insoluble drugs such as TAM and that encourage mice to voluntary consume the drug, while retaining efficient TAM mediated induction of gene expression. Our results show that formulations in which TAM is first dissolved in oil and then dispersed in sweetened milk or syrup encourage mice to voluntarily consume TAM offered with a micropipette. The efficacy of the new formulations, as reflected by TAM-mediated genetic recombination, was tested in two CreER^T2^-based transgenic mouse models. First-pass metabolism was assessed by comparing serum concentrations of TAM-metabolites from treated mice using a cell-based *in vitro* assay. Together, these results demonstrate that feeding TAM-sweetened-milk formulations is a more animal-friendly administration method to replace OG and therefore could become the method of choice when administering water insoluble drugs.

## Results

### Mixing oil with sweetened solutions makes it palatable to mice

To adapt the micropipette-guided drug administration to tamoxifen, we first compared the palatability of TAM dissolved in oil before and after adding sweetened condensed milk or berry syrup. To encourage voluntary consumption and drinking from a micropipette tip ^7, 8, 10^, mice were trained for three days with the different formulations before being offered the same formulation containing TAM on the fourth day (Fig. 1A). The palatability of the solutions was assessed by recording the time the mice needed to consume a specific volume ^32^. We defined consumption as voluntary when mice drank the substance within 60 s while sitting on the cage grid. To prevent the mice from wandering or leaving the grid, they were gently held by the tail (Fig. 1B and detailed in the methods). When offered peanut oil (Fig. 1D, first panel) or corn oil (not shown), vehicles typically used to dissolve and administer TAM via OG, almost none of the mice voluntarily drank the oil during training or when subsequently offered oil-containing TAM.

**Figure 1:**
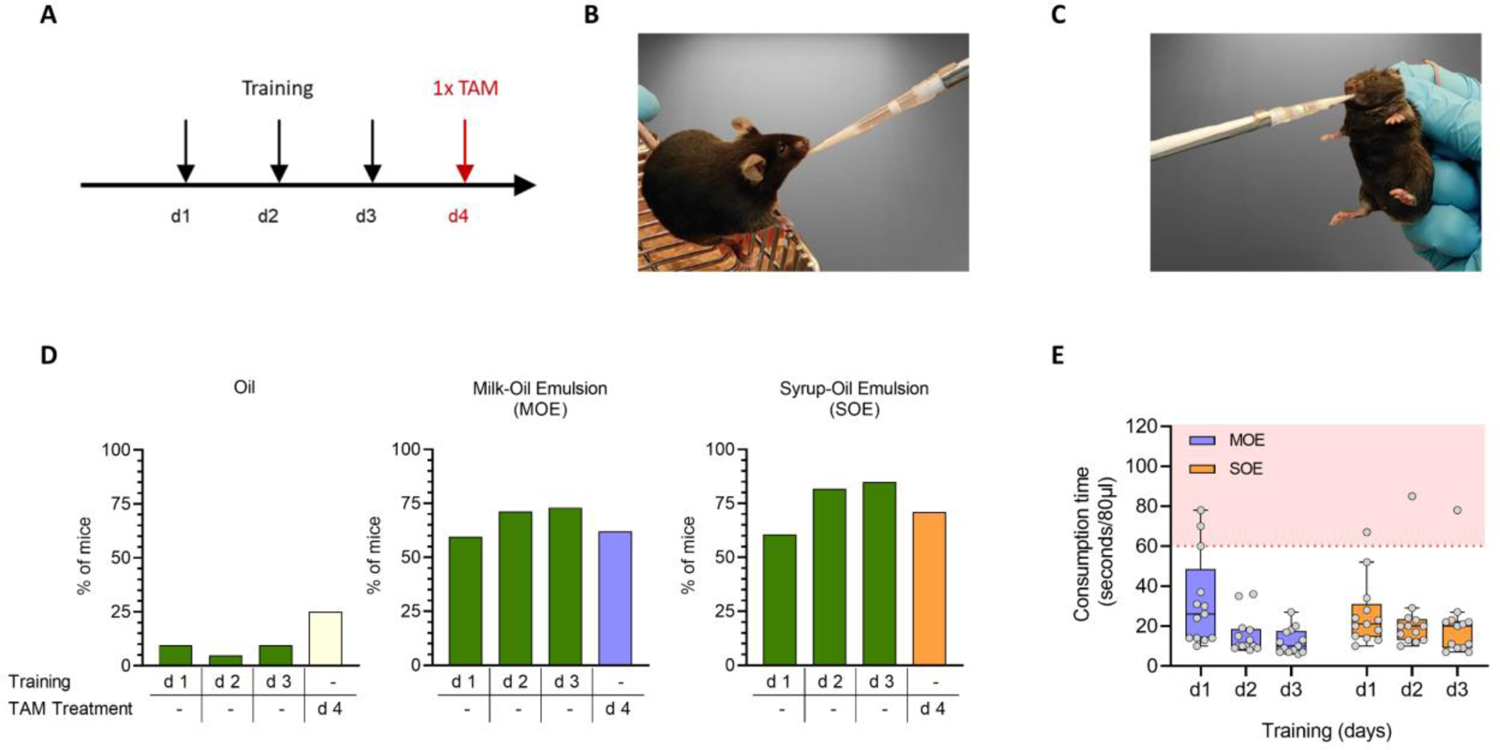
Sweetened oil emulsions and training improves voluntary drug consumption. (A) Mice were trained for three days to consume 80 µl of formulations from a disposable plastic micropipette tip and on the fourth day were offered the same formulation containing TAM. (B) Feeding time was recorded and considered voluntary if the mouse, held slightly by the tail, drank the offered volume in less than 60 s. (C) Any formulation not consumed after 60 s was offered again after gently restraining the mice by the scruff, and the extra time required to drink the remaining solution was recorded. (D) Fraction of mice that voluntarily consumed (<60 s), oil (Oil, n=21), sweetened milk-oil-emulsion (MOE, n=52), or syrup-oil-emulsions (SOE, n=33) during three days of training (green bars) or TAM containing formulations on the fourth day. Yellow bar = OIL-TAM (n=12), blue bar = MOE-TAM (n=17), orange bar = SOE-TAM (n=17). The oil used for these formulations was sterilized by heat treatment. (E) Total consumption time for three training days, recorded from the moment the MOE or SOE formulation were offered (n=13 per group). Indicated is the time before restraining (white zone) as described in (B) and after restraining (red zone) as described in (C). The oil used for the emulsions was sterilized by filtration. Minimum and maximum values, interquartile range, and median are depicted as Tukey boxplots with individual data points shown as grey circles.

The addition of sweetened milk has been shown to encourage mice to voluntarily consume water-soluble drugs^7, 8^. However, simply mixing oil with sweetened condensed milk to improve palatability generated solutions that quickly coalesced (data not shown), regardless of how long the solutions were mixed with an electric vortex mixer. In contrast, using a simple high energy homogenization method, the so-called 2-syringe method^33^ (Fig. S1A and described in methods), resulted in stable oil-in-water emulsions. We retained two promising formulations, a milk-oil-emulsion (MOE) made with sweetened condensed milk and a syrup-oil-emulsion (SOE) made with berry syrup. The mean diameter of the oil droplets in these emulsions as determined by microscopy (Fig. S1B) was 13±4 µm (n=402) for MOE and 15±5 µm (n=436) for SOE.

To ensure accurate drug administration, we verified that the emulsions can be dispensed precisely and consistently using a micropipette. This was confirmed by the low coefficients of variation observed when a fixed volume was repeatedly pipetted. (Fig. S1C).

Both MOE and SOE emulsions improved voluntary consumption of oil, with more than 60% of the mice drinking the sweetened solutions within the first 60 s, while only 10% of the mice voluntarily consumed oil without any additives. After three days of training, the fraction of animals that voluntarily consumed the offered solutions increased to more than 75% for MOE and SOE, while it remained unchanged for oil (Fig. 1D). Training with the sweetened emulsions shortened the manipulation time for most mice to just 10 to 30 s per administration (Fig. 1E and Fig. S2A). Since it takes some time for the mice to become aware of the pipette tip, the actual drinking time is even shorter. Training is thus important since it results in shorter handling times, which is beneficial for the mice and minimizes overall experimental procedure times.

### Sweetened emulsions and training encourage TAM consumption

To test whether the sweetened emulsions also improved TAM consumption, mice were trained for three days with MOE or SOE and on the fourth day received MOE or SOE containing TAM at a final dose of 80 mg/kg. Similarly, control mice were pipette-fed with oil (training) and then with OIL-TAM (80 mg/kg). We observed that 75% and 60% of the mice voluntarily consumed MOE-TAM and SOE-TAM, respectively, versus 25% for OIL-TAM solutions (Fig. 1D). Interestingly, when the oil was sterilized by filtration instead of heat treatment, the mice more readily consumed the sweetened emulsions (Fig. S2A), suggesting that heating the oil affected the taste of the emulsions. Indeed, with filtered oil, the fraction of mice that voluntarily consumed the emulsions during training and subsequent administration of TAM increased to 100% (Fig. 1E and Fig. S2A-C).

The use of transparent tunnels in husbandry and experiments has been described as beneficial to animal welfare ^34^. Therefore, we incorporated tunnel handling into our procedures. Although we found that the tunnels were helpful for removal from the home cage, weighing, identification, and transfer to a cage grid for feeding (Fig. S4), the animals could not be motivated to drink the emulsions from within a tunnel.

Taken together, the use of sweetened oil-in-water emulsions motivates mice to voluntarily consume TAM after at least two days of training.

### Efficacy of pipette-administered MOE-TAM is non-inferior to that of gavaged OIL-TAM

Having shown that sweetened emulsions encourage voluntary consumption of TAM, we next compared the efficacy of TAM administered with a micropipette using SOE and MOE formulations, with that obtained after oral gavage of OIL-TAM. The capacity to induce Cre-mediated recombination was analyzed using a mouse model where the CD4-CreER^T2^ knock-in strain^35, 36^ is crossed with mice containing an inserted transgene, in which the VαJα exon of the alpha chain of an HY-specific T cell receptor (TCRα) is flipped into transcriptional orientation following Cre-mediated recombination. In the correct orientation, the complete TCRαβ is expressed on the cell surface of thymocytes due to the presence of a conventional transgene in these mice coding for the corresponding HY-TCRβ chain^37^.

Using these so-called HY-switch mice, thus mice that will express the HY-specific TCR on thymocytes when treated with TAM, we performed a non-inferiority analysis to determine if the new formulations are not worse than, or ‘non-inferior to’, the standard OG treatment^38^. The experiment was prospectively powered (see methods) based on preliminary data from HY-TCR expression induced after conventional TAM treatment (oral gavage of OIL-TAM). Induction of HY-TCR expression (% HY-TCR positive cells) on thymocytes was analyzed 40 hours after pipette-feeding the mice with a single dose of MOE-TAM, SOE-TAM or, as control, OIL-TAM administered via OG. With each treatment, mice received 80 mg of TAM per kilogram of body weight. For the emulsions (MOE-TAM and SOE-TAM) such an accurate and mouse-tailored dose administration was achieved simply by changing the volume dispensed by the micropipette. However, a similar accurate dosing is not possible with the syringes that are typically connected to the gavage needle. Since accurate dosing was essential to correctly compare the efficacy of the different methods, we prepared weight-adjusted OIL-TAM solutions for each mouse in the OG group, of which the corresponding mouse then received 200 µl.

The efficacy of the different TAM formulations in inducing HY-TCR gene expression was quantified by flow cytometry. The results of three independent experiments were combined to achieve the number of mice per group (n=13) required for non-inferiority testing (with each experiment comparing equivalent group sizes). This data showed that pipette-fed MOE-TAM but not SOE-TAM is non-inferior to standard OG-OIL-TAM. Indeed, the 95% confidence interval (CI) of the difference between the mean of reporter (HY-TCR) induction in the OG-OIL-TAM group and that of the MOE-TAM treated group is located within the noninferiority zone (Fig. 2A). This is also reflected by the mean TCR induction observed in the MOE-TAM group which is similar to that of the OG-OIL-TAM group (47% vs 48%) (Fig. 2B), in contrast to the SOE-TAM group (31%). As a negative control, mice were trained for three days with MOE but not treated with TAM (n=6). In these mice, as well as in untreated mice (data not shown), HY-TCR expression in the thymus is undetectable (Fig. 2B).

**Figure 2:**
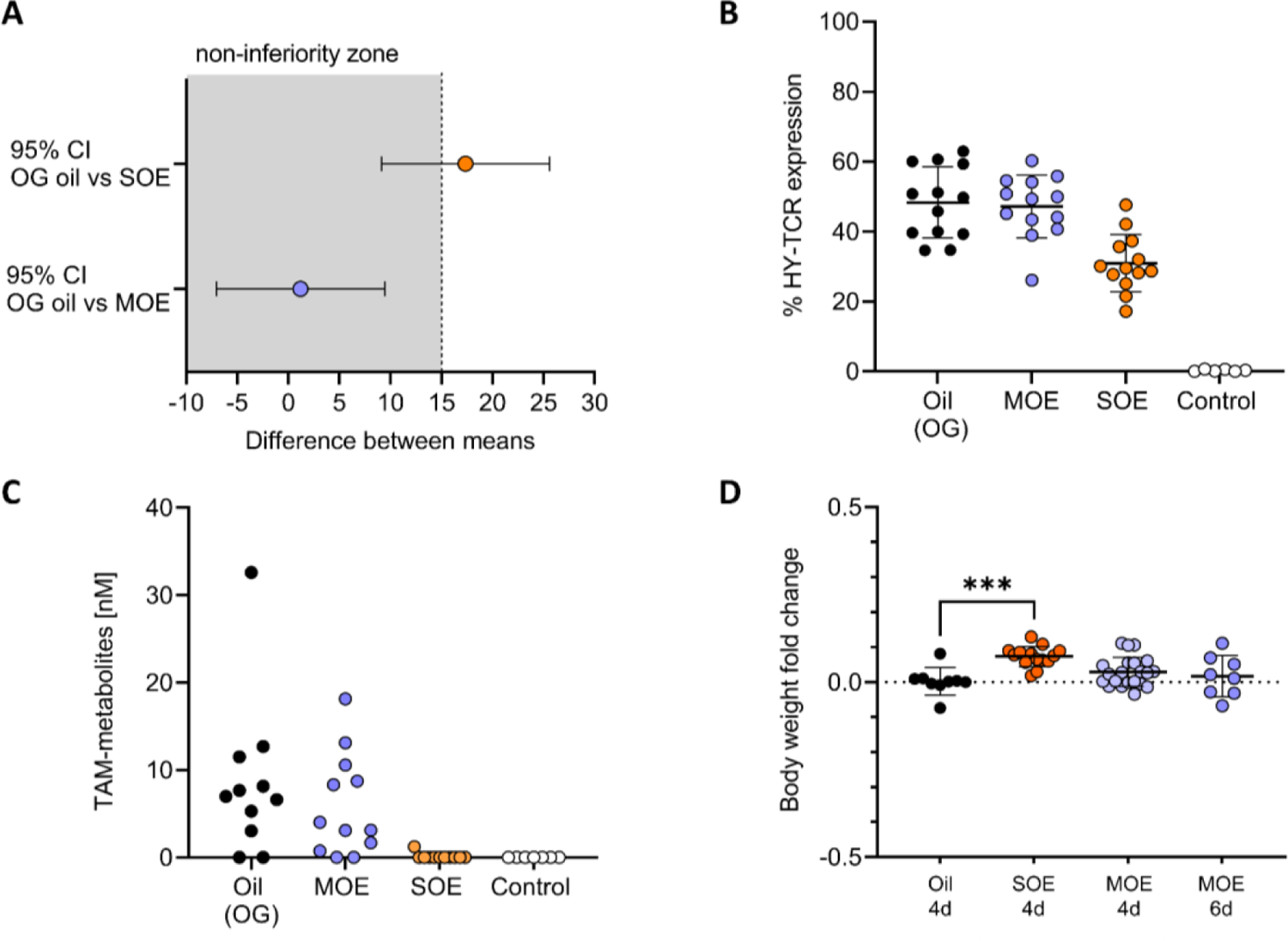
Efficacy of tamoxifen-induced gene expression. Comparison of CreER^T2^-dependent HY-TCR reporter expression in thymocytes of female mice, 40 hours after treatment with a single dose of 80 mg/kg TAM, administered in oil via oral gavage (OG) or in oil-in-water emulsions made with sweetened milk (MOE) or syrup (SOE) using the micropipette method. Controls were HY-TCR knock-in mice, trained (3 d) with MOE but not treated with TAM (n=6). (A) Non-inferiority graph, depicting the 95% confidence interval (CI) of the difference between the mean of the OG oil group and SOE group or MOE group of the results shown in panel B. The 95% CIs were computed using Dunnett’s confidence interval formula following the analysis of variance. The non-inferiority margin (d) was set at 0.18 (1.5-times SD of control OG-oil group). (B) The percentage of HY-TCR expressing thymocytes was determined via flow cytometry as described in the methods. Shown are means ± SD per treatment group (n=13 per group). (C) TAM-metabolite concentrations in the serum of mice described in A, were assessed in vitro using MEFs from R26-Cre-ER^T2^/Ai14 mice. (D) Fold change in body weights relative to the oil group following daily administration of the indicated formulations for four or six days. Group sizes: Oil n=9, SOE n=14, MOE n=21. Depicted are means ± SD. *** p=0.0006 according to one-way ANOVA followed by post-hoc Dunnett’s multiple comparison test.

Interestingly, SOE but not MOE treated groups significantly gained weight compared to the control group receiving oil only (Fig. 2D). These results thus show that only MOE-TAM is equally efficient in inducing CreER^T2^ activation and offers an animal-friendly alternative to OG for the administration of TAM. An additional advantage of the micropipette administration method is that it allows for accurate dosing of TAM without the need to make separate OIL-TAM solutions for each mouse, thereby also improving experimental conditions and reproducibility.

### Serum concentrations of TAM metabolites correlate with CreER^T2^-induced gene expression

CreER^T2^ activation in mice after TAM treatment is primarily mediated by its major bioactive metabolite 4-OH TAM^39, 40^, since TAM itself is a poor inducer of CreER^T2^ ^39, 41, 42^. Thus, Cre-mediated recombination is highly dependent on efficient first-pass metabolism of TAM in the liver^22, 39, 40, 43^. To directly assess serum concentrations of bioactive TAM metabolites in the different treated mice as a measure of first-pass TAM metabolism and therefore compare the pharmacological properties of the different formulations, we developed an in vitro assay based on mesenchymal embryonic fibroblasts (MEFs) isolated from R26-CreER^T2^/Ai14 mice. When these MEFs are exposed to 4-OH TAM, a constitutively expressed recombinant CreER^T2^ protein mediates the removal of a loxP-flanked STOP cassette, resulting in the expression of a red fluorescent tdTomato protein. As shown by time-lapse microscopy and flow cytometry of tdTomato expression (Fig. S3A-C), induction in these cells is time and dose-dependent.

To test if these MEFs can be used to quantify serum TAM metabolites, we compared serum metabolite concentrations obtained with R26-Cre-ERT2/Ai14 MEF to those obtained with an EU-approved assay^44^, known as the LUMI-CELL^®^ ER assay^45^. This latter cell culture-based assay was developed for the quantification of estrogen receptor (ER) agonists^45, 46^ but it can also detect ER antagonists such as 4-OH TAM^46^. We used sera collected from C57BL/6 mice, 6 h after OG treatment with OIL-TAM (40 or 80 mg/kg) and compared the results obtained with both assays. Although the principle of TAM metabolite detection is different, both assays gave comparable concentrations for each serum (Fig. S3D), confirming that the R26-CreERT2/Ai14 MEF can indeed be used to determine and compare serum metabolite concentrations. Since reporter expression in the MEF directly measures CreER^T2^ activation, we used this assay to compare bioactive serum TAM metabolite concentrations from CreER^T2^ mice that were treated with the different TAM formulations.

In line with the induction of HY-TCR expression (Fig 2A-B), treatment with OIL-TAM by OG or pipette-feeding with MOE-TAM resulted in comparable serum concentrations of TAM metabolites (Fig. 2C), confirming that first-pass metabolism of TAM administered in sweetened milk emulsions is comparable to that obtained with conventional OG. Serum TAM metabolite concentrations typically peak at 6-7 hours after TAM administration and then rapidly decrease over the next 24 to 48 hours^47^. Accordingly, at the time-point when the mice were sacrificed, which was optimal to evaluate gene expression, only low metabolite concentrations (nM range) were observed (Fig. 2C). None of the mice fed with SOE-TAM had detectable serum TAM metabolites at the time of analysis, in line with the lower levels of TCR induction observed with this formulation.

Together, our results indicate that voluntary consumption of MOE-TAM formulations is as efficient as conventional OG treatments regarding reporter gene induction and generation of serum TAM metabolites. In contrast, TAM in syrup emulsions is less effective compared to both OIL-TAM (OG) and MOE-TAM formulations and in addition results in significant gains in body weight.

Thus, while both sweetened milk and syrup-based emulsions encourage mice to voluntarily consume TAM offered with a micropipette, only the sweetened milk-oil emulsions offer an acceptable alternative method to OG.

### Repeated administration of MOE-TAM increases the induction of reporter gene expression

In the experiments described above, HY-switch mice were treated with only one dose of TAM to induce a cohort of TCR expressing cells. While such one-time treatment is also used for other mouse models^48^, most in vivo studies involving CreERT or CreER^T2^-mediated gene expression require repeated treatments to ensure sufficient genetic recombination in target tissues or cells^11, 35^. Therefore, we tested the efficacy of treating animals with MOE-TAM once every 24 hours for a total of five consecutive days, a treatment regimen recommended for many CreER^T2^-based mouse models^11, 49^. As observed before, the mice voluntarily consumed MOE-TAM on the first day. However, they were reluctant to readily consume a second dose the next day and instead required restraint after 60 s to finish drinking the offered volume (Fig. 3A). Since split dosage treatment of mice, twice a day, was shown to result in similar recombination frequencies as a full TAM dose^50^, we tested whether this apparent aversion could be overcome by halving the TAM dose to 40 mg/kg. However, most mice also refused to voluntarily consume a second serving of this lower amount, although they had readily drunk the first dose (Fig. 3B). Interestingly, once the mice were restrained (i.e., after 60 s) they quickly drank the formulation (Fig. 3A & B), despite having shown little or no interest in it while sitting on the grid. Therefore, we tested whether mice would also rapidly consume additional treatments if they were immediately held by the scruff. Indeed, mice consumed additional treatments without hesitation while being held by the scruff (Figure 3A-D) and this independently of the dose offered (40 mg/kg or 80mg/kg). Therefore, feeding multiple doses of MOE-TAM is possible, but is best performed by gently restraining the mice from the second dose onwards.

**Figure 3:**
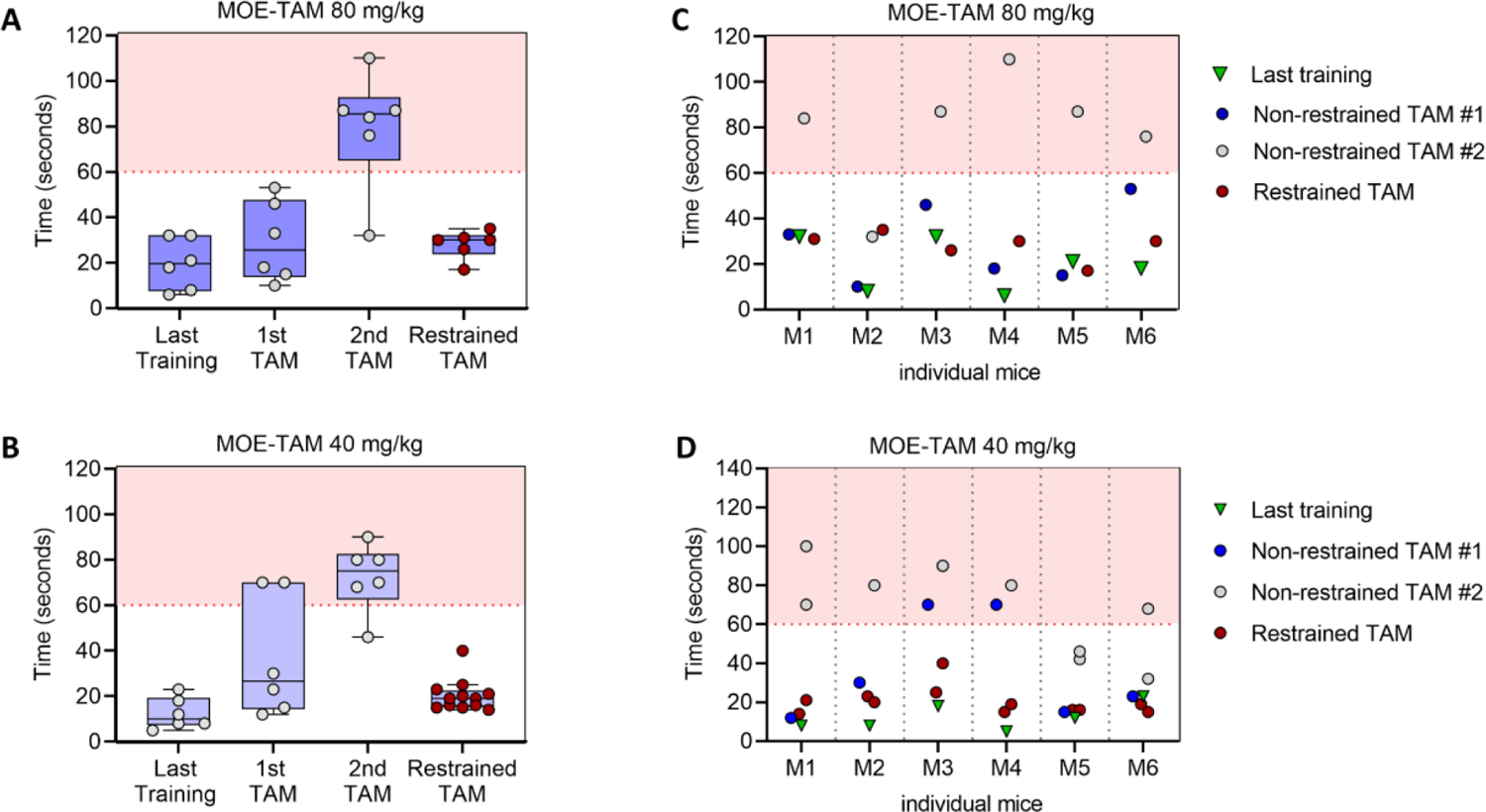
Repeated administration of TAM using milk-oil emulsions (MOE). Mice were trained for three days before MOE-TAM treatment. Consumption time for the last training, first and second TAM treatment was recorded and displayed as described in Figure 1E. For subsequent MOE-TAM treatments, mice were immediately restrained (red dot plots). (A) and (C) Treatment with 80 mg/kg TAM in MOE. (B) and (D) Treatment with 40mg/kg TAM in MOE. (C) and (D) Consumption time for individual mice (M1 to M6) for data shown in (A) and (B). Mice were either offered the formulations without being restrained for the first 60 seconds (grey or blue dot plots) or immediately restrained (red dot plots). For (A) and (B) minimum and maximum values, interquartile range and median are depicted as Tukey boxplots with individual data points shown as scatter dot plots.

Since mice readily consume sweetened MOE-TAM emulsions for at least five days, we evaluated the efficacy of such repeated MOE-TAM treatments using R26-CreER^T2^/Ai14 reporter mice, the same mouse line that was used to derive the R26-Cre-ERT2/Ai14 MEFs for the in vitro assay. This mouse strain is more suitable for monitoring long-term TAM treatment due to the ubiquitous and additive expression of the TdTomato reporter. We compared two groups of mice, namely animals treated daily with MOE-TAM for five days and a second group that received MOE-TAM only on day 5 (Fig. 4A). The expression of TdTomato was evaluated by flow cytometry in thymocytes and splenocytes (Fig. 4B) and by fluorescence microscopy of spleen sections (Fig. 4C). These results show increased induction of tdTomato reporter expression after repeated feeding of MOE-TAM compared to one-time treated mice, reflecting efficient repeated delivery of TAM and increased induction of CreER^T2^ activation.

**Figure 4.**
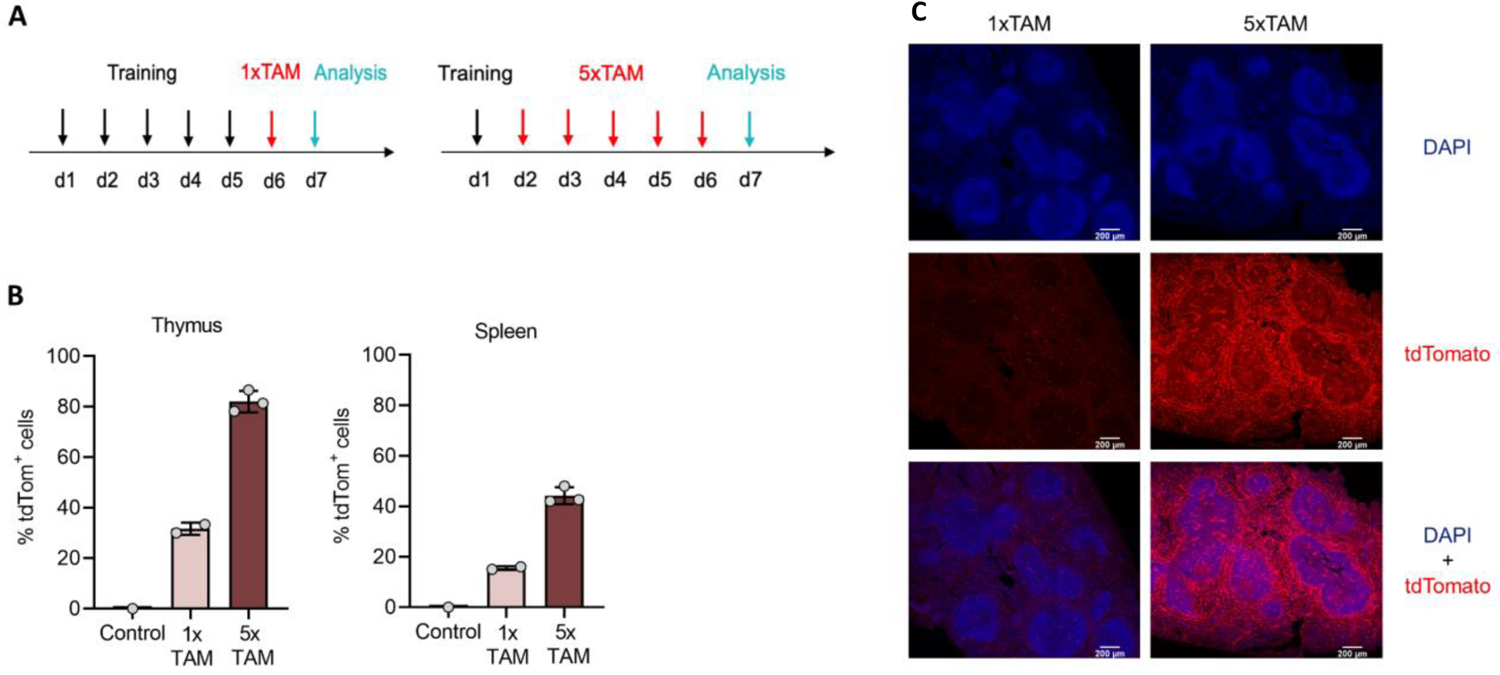
Multiple administration of TAM in milk-oil emulsions (MOE). (A) Induction of tdTomato expression was assessed in R26-Cre-ER^T2^ Ai14 tdTomato mice after training and TAM treatments as indicated. Mice were either offered MOE-TAM (80 mg/kg) once a day for five days (n=3) or only one day (n=2) or did not receive TAM (n=1). (B) The percentages of thymocytes and splenocytes expressing tdTomato were determined by flow cytometric analysis. Bar graphs depict means ± SD. (C) Representative microscopy pictures of tissue sections from spleens collected from five times or once treated mice. Tissue sections were stained with DAPI (blue) and analyzed for tdTomato protein expression (red). Magnification 20x and scale bar=200 μm. All images were processed using the identical settings for each channel.

In conclusion, our results show that repeated administration of MOE-TAM results in greater reporter induction. However, such treatments are best performed by holding the mice by the scruff, except during training and first TAM treatment.

## Discussion

Animal studies are important for the advancement of our understanding of animal and human physiology, as well as for the development of new therapies^51–53^. However, some current laboratory practices, even when used for years as standard procedures, are suboptimal in terms of animal welfare. Physiological changes associated with discomfort or stress have been reported to interfere with and confound experimental outcomes^24–27^, potentially compromising the reliability and reproducibility of experimental results. Over the past few years, new procedures have been explored to reduce stress and injuries related to drug administration. Such methods include voluntary drug feeding of rodents using palatable vehicles (e.g., sucrose water, peanut butter, tablets, jam, or jelly)^54–57^. Unfortunately, these methods do not allow accurate drug delivery, potentially resulting in over or underdosing^10, 58^, or require individual housing which was shown to inflict stress on animals^59^.

A promising drug administration method that allows accurate drug delivery with minimal or no induced stress responses is the so-called micropipette-guided drug administration^7,8^. Although this method was shown to replace IP and OG with similar efficacy for the drugs tested, up until now it was only reported for the administration of water-soluble drugs. In the current study, we show that mice can be encouraged to voluntarily consume water-insoluble drugs such as tamoxifen using a micropipette by administering the drug in sweetened ‘oil-in-water’ emulsions. The mice eagerly consumed both sweetened milk- and syrup-based emulsions, but only feeding TAM in sweetened milk emulsions (MOE-TAM) resulted in CreER^T2^-mediated reporter expression that is statistically non-inferior to OG of OIL-TAM solutions. Stable oil-in-milk emulsions were obtained by a simple high-energy homogenization method using two connected syringes^33^, which is cost-effective and readily available to most research groups. Although we did not extensively explore alternative methods, more expensive professional homogenizers could allow more automated processing^60^. Larger volumes of MOE can be prepared for training and be frozen and thawed without coalescing (not shown). However, this may not be possible for emulsions containing TAM, since TAM readily precipitates at low temperatures.

Based on our observations, training for at least 2 days is indispensable for efficient consumption of MOE-TAM solutions, possibly because rodents are neophobic towards new foods^61, 62^ and need to be first habituated to drinking from the pipette tip and/or to the taste of the formulations. Sterilizing the oil by filtration instead of heat treatment further improved consumption of the emulsions, suggesting that changes in taste can affect palatability. The method is also affected by the preferences of individual mice (Fig. 3C & 3D), as reflected by differences in consumption times. Long-term daily treatments using pipette feeding of sweetened formulations were reported for water-soluble drugs^7,8^. We show that repeated daily pipet feeding of MOE-TAM is also possible, albeit only when the animals are gently held by the scruff after the first dose of TAM, suggesting the mice acquired a mild taste aversion ^54, 63^. Although we did not explore longer treatments than 5 days, we expect that additional daily administrations with MOE-TAM will also be possible.

Whereas mice readily accepted pipette feeding of both SOE-TAM and MOE-TAM, the reduced serum concentrations of TAM metabolites and gene induction in mice treated with SOE-TAM suggest that substances in syrup affected the efficacy of TAM. In contrast, treating mice with TAM in sweetened milk emulsions (MOE-TAM) resulted in CreER^T2^-mediated gene expression that is equivalent to conventional OG of OIL-TAM, indicating that the administration of TAM and first-pass metabolism are unaffected by sweetened milk. Administering TAM in MOE formulations for at least 5 days, which we confirm to result in enhanced reporter gene recombination and expression, will allow to successfully apply this method to a larger number of different CreERT2 mouse models.

The administration of drugs, including TAM dissolved in oil, by pipetting directly into the mouth of mice has been previously reported^64, 65^. Though these methods do not rely on palatable drug formulations or voluntary consumption by mice. Instead, the mice were forced to swallow the solution by pressing the pipette tip against the hard palate^64^ or behind the diastema of the mouth^65^. While this method is likely to be more stressful than our voluntary administration method, efficient reporter gene induction was reported when TAM was administered in this manner^65^. However, TAM delivery appears to be suboptimal because consistent results were only obtained with very high TAM doses (7.5-10 mg/mouse corresponding to 300-400 mg/kg)^65^. Differences in CreER^T2^ strains used in that study as compared to our experiments could explain the requirement for different TAM doses. However, in this previous study, no direct comparison was made with conventional OG or IP administrations^65^, and neither was it shown or discussed if mice can be treated with more than one dose with this method. In contrast, in our study we demonstrate that the efficacy of palatable MOE formulations is statistically non-inferior to that obtained after OG of identical TAM doses. Furthermore, we obtain consistent induction of the HY-TCR reporter within groups of mice, even with MOE-TAM doses as little as 0.5 mg TAM per mouse (20 mg/kg) (data not shown). In our experience, any formulation that is not actively licked up by the mice will result in loss of material, leading to variable doses and thus variable experimental outcomes. We observed that feeding oil only is very inefficient. Drops of oil tend to rapidly spread in the fur around the mouth of the mice and most mice just kept the oil in their mouth without drinking it, suggesting that consumption cannot be forced, even when the mice are held by the scruff.

Although treatment with palatable formulations present many advantages, there are also some limitations to consider. The respective drug could interact with the vehicle, thereby affecting efficacy of the treatment, as observed for syrup and TAM (SOE-TAM) in our experiments. Depending on the drug tested and the experimental read-outs, the sweetening substances in the formulations could compound experimental outcomes. Additionally, the vehicle itself may contain activities relevant to the measured variable. Finally, pipette feeding may not work when a drug cannot be made palatable by sweetening, for example if the drug itself has a strong unpleasant taste. For such drugs or those that induce strong adverse effects, oral gavage might still be necessary. We observed that the mice showed a mild aversion after a first treatment with MOE-TAM, requiring holding the mice by the scruff for feeding additional daily dosages. Such gentle restraint could interfere, for example, with behavioral studies or induce some level of stress, potentially affecting experimental results. However, because the mice still voluntarily consumed TAM, pipette feeding is still preferable to alternatives such as OG or IP injections which also require restraint.

Alternative methods to prepare TAM-containing micro-emulsions have been reported as replacements for tablets to treat breast cancer patients. However, these micro-emulsions did not contain sweet substances, were made with detergent-based emulsifiers, and were either tested solely in vitro^66^ or were administered to tumor-bearing mice via oral gavage^67^.

There is little consensus in the scientific community on the TAM dosage necessary to induce sufficient CreER^T2^-activation in mice^12, 50^. The reasons may lie in variable exposure of different target tissues, differences in locus accessibility for recombination, kinetics in gene expression, and finally protein stability. In most published experiments, typically all mice receive the same volume of TAM through OG regardless of body weight, prompting the preferential use of high dosages to ensure complete recombination. While administered volumes can be adjusted to some extent with the syringes used for OG, micropipette administration of MOE-TAM formulations allows for more accurate treatments over a wide range of dosages by adjusting both the administered volume, as well as the concentration of TAM in the sweetened milk emulsions. In our experience, 80 mg/ml of TAM in oil, therefore a final 40 mg/ml of TAM in the MOE emulsion, is the recommended maximum concentration to use. Higher TAM concentrations could result in precipitation of TAM in the emulsions, especially below room temperature, which will not only affect accuracy of treatment, but could also cause adverse effects in mice^68^. High TAM concentrations such as 200 mg/kg are in any case not recommended as they do not increase the rate of recombination, but rather can lead to metabolic stress and increased lethality^50^. Using the pipette administration method, we successfully administered TAM from 20 mg/kg to 120 mg/kg with MOE-TAM volumes ranging from 30 µl to 120 µl using a 200 µl micropipette, including mice as young as five weeks (data not shown).

Administration of TAM by chow or drinking water has been used for some experimental applications with the disadvantage that the timing and amount of drug consumption are not controlled^10, 58^, mice reduce their food or water intake^54, 58^, or required single-housing of the animals^5, 69–71^. Single housing of rodents is a known stressor with physiological consequences^27, 72^ and is thus strongly discouraged.

Several studies have suggested that a more humane approach to reducing pain and distress experienced by laboratory animals will have a positive impact on behavioral and physiological processes and therefore reduce variability in experimental data^24–27^. Thus, reducing pain and distress will not only benefit animal welfare but also, because of reduced experimental variability, will help conduct more reliable and robust experiments and reduce the number of experimental animals required per experiment.

Taken together, dosing with the micropipette method is a valuable, more animal-friendly alternative to more invasive methods such as OG and IP injections. Because administered volumes can be easily adjusted with the micropipette, dosing is also more accurate, and holding the mice by the tail or the scruff can be performed with little training of the experimenter.

We used TAM as an archetype drug to implement and evaluate a non-invasive administration procedure for water-insoluble drugs based on voluntary consumption. This new method adds to a growing number of applications that aim to replace more invasive administration routes, such as OG or injections, with animal-friendly methods that are equally accurate in dosing, timing, and outcome^7, 8^.

## Methods

### Animals

All mouse strains used in this project were of C57BL/6J genetic background. HY-switch mice were obtained by crossing the following strains: CD4-CreER^T2^(B6.CD4^tm1(Cre/ERT2)ThBu^)^35^, HYβtg mice (B6.Tg(Tcrb)93Vbo)^37^ and HYasn (B6.TCRA^tm2ThBu^ (unpublished)). The resulting HY-switch mice express the CreER^T2^ protein under the control of the CD4 promoter and have a loxP-flanked HY-TCR VαJα exon inserted into the TCRα locus so that expression of HY-TCR is only possible after Cre-mediated inversion and the presence of the HY-TCRbeta transgene (HYβtg). The TCR δ locus was deleted by Cre-mediated recombination in the germline. Only female HY-switch mice (10-18 weeks of age) were used for the non-inferiority study since HY-TCR expressing thymocytes are specific for the male HY self-antigen and are readily eliminated in male mice due to negative selection^73, 74^. The tdTomato reporter mice used in this study were generated by crossing R26-CreER^T2^ mice (B6.129-Gt(ROSA)26Sor^tm1(cre/ERT2)Tyj^/J)^75^ and Ai14 mice (B6.Cg-Gt(ROSA)26Sor^tm^^14(CAG-tdTomato)Hze^/J)^76^. Both males and females between 10-18 weeks of age were used. All mice were bred in-house under SPF conditions in the absence of FELASA-listed pathogens and group-housed in individually ventilated cages in rooms with controlled temperature, humidity, and light cycles. All mice had *ad libitum* access to food and water throughout the study. Mice were randomly assigned to a treatment group and group-housed accordingly. Mice were acclimated for at least seven days before experimental use. All experimental procedures described were approved by the Cantonal Veterinarian’s Office of Zürich (License number ZH107/2020).

### In vitro TAM assays

TAM metabolite detection was performed with the experimenter blinded to the serum samples. The LUMI-CELL® ER assay is based on the VM7Luc4E2 cell line, generated by Prof. M. S. Denison (University of California – Davis) by transfection of human MCF7 breast cancer cells with the pGudLuc7 plasmid^45, 77^. The luciferase (Luc) reporter in VM7Luc4E2 cells responds in a dose-dependent manner to estrogen receptor (ER) agonists or antagonists (eg. 4-OH TAM)^44, 45, 77^. VM7Luc4E2 cells were cultured in estrogen-free medium as published^45, 78^. Estrogen-depleted VM7Luc4E2 cells were seeded at 6.6×10E4 cells/well in 100 µl assay medium in 96-well plates and cultured at 37 °C with 5% CO2. Twenty-two hours later, per well, 100 µl of media were added containing estrogen (0.1 nM), 4-OH TAM, or diluted mouse sera (final concentration 15%). Each sample was tested in triplicate. After 22h, cells were washed twice with DPBS, and luciferase activity was recorded using the Promega Luciferase assay system (Promega #E4030) according to the manufacturer’s instructions and as published^45^ using a TECAN reader (SPARK®TECAN). The standard curve of inhibition of estrogen-induced Luc expression in response to serial dilutions of 4-OH TAM (5-80nM) was fitted by a sigmoidal curve (R2 > 0.9) according to the Prism 4-parameter fit algorithm (4PL) and used to calculate the concentration of TAM metabolites for each tested mouse serum (Prism v9.2.0, GraphPad).

For the CreER^T2^-Ai14 reporter assay, mesenchymal embryonic fibroblasts (MEF) were isolated from individual R26-CreER^T2^-Ai14 mice and cultured as previously described^79^. Selected clones were immortalized by transfection with the plasmid pBSSVD2005, coding for the large T oncogene SV40 (V40 1: pBSSVD2005 was a gift from David Ron; Addgene plasmid #21826; http://n2t.net/addgene:21826; RRID:Addgene_21826), using LipofectamineTM LTX (Invitrogen #L3000-008) according to the manufacturer’s instructions and selected using a protocol from Heather P. Harding^80^. To detect serum TAM metabolites, MEF were seeded in 24-well plates (Falcon #353047) at 5×10E4 cells per well in 300 µl growing medium and cultured at 37°C with 5% CO2. After 4-5 hours, 100 µl of diluted mouse serum (final 15%) or serial dilutions of 4-OH TAM (1 nM - 30 nM 4-OH TAM) were added to duplicate wells. At the time points indicated in the figures, cells were washed and detached with trypsin and tdTomato expression measured with a BD LSR Fortessa II (BD). Results were analyzed using the FlowJo™ v10.4 Software (BD Life Sciences). The percentage of tdTomato-positive cells in each well was determined by manual gating, as indicated in Fig. S3B. The standard curve of responses (average of duplicate wells) to serial dilutions of 4-OH TAM, was fitted by a sigmoidal curve (R2 > 0.9) according to the Prism 4-parameter fit algorithm (4PL) and used to calculate the relative concentration of metabolites of TAM for each mouse serum (Prism v9.2.0 GraphPad).

For time lapse microscopy, MEFs were seeded in 8-well imaging chambers (Ibidi #80826) and treated with 7nM 4-OH TAM, 70nM 4-OH TAM, or left untreated. Selected fields (8 for each well) were imaged using an inverted widefield microscope (Zeiss, Axio Observer Z1, Germany) provided with temperature and CO_2_ control (37°C and 5% CO_2_), every 20 min for a 23 h period with a 63x objective. Images were processed and analyzed using ImageJ software (v1.53c National Institutes of Health, MD, USA).

### Preparation of Emulsions

Peanut oil (Sigma #P2144) was sterilized at 160°C for 3h or by filtering (0,22 µm Steriflip filter, Sigma #C3238). TAM (Sigma #T5648) was dissolved in 100% ethanol (Avantor #32221) at 100 mg/mL and mixed with an identical volume of peanut oil^35^. The oil/ethanol/TAM solution was heated in a sonicator bath (Transsonic Digital D-7700 Elma) to 58°C. A vacuum was applied to accelerate the evaporation of the ethanol. For each experiment, a fresh 100mg/mL TAM in oil stock solution was prepared and used to make the different formulations as indicated.

Milk-oil-emulsions (MOE) for this study were prepared with sweetened condensed milk (Nestlé Milchmädchen - Sweetened Condensed Milk with 54.7% sugar, 8.5% fat, 20.5% non-fat milk dry mass) diluted 1:2 with sterile double distilled water (Ecotainer, Braun, #82479E-E). Other brands of condensed milk (e.g., MIGROS Kondensmilch™, Migros, Zurich, Switzerland) are also suitable^7,8^. MOE emulsions were made using one volume of peanut oil and one volume of diluted milk, homogenized using the two-syringe method (also known as syringe extrusion) as published^33, 81^ and detailed below. MOE-TAM emulsions were prepared with oil containing 50 mg/ml TAM (for treatments with 80 mg/kg) or 25 mg/ml TAM (for treatments with 40 mg/kg).

Syrup-oil-emulsions (SOE) were prepared using two volumes of undiluted syrup (Coop, Switzerland, with 81% sugar, 40% berry juice from concentrate, acidifier E330, H2O, and flavoring) and one volume of peanut oil. SOE-TAM emulsions were made as above but with oil containing 75 mg/ml TAM (for treatments with 80 mg/kg).

All reagents and solutions were kept sterile. Both MOE and SOE emulsions were freshly made on the day of treatment by emulsification during 10 minutes at room temperature using 2 mL Luer-Lock syringes (Braun # 4606701V) connected with a one-way Luer female to female adaptor (Cadence Science #6521IND).

Oil and emulsions were always dispensed using reverse pipetting^82^. The accuracy of pipetting emulsions was tested by repeated dispensing and weighing (microbalance) of a fixed volume (75 µl).

For determining the diameter of the oil droplets in the emulsions, the fluorescent dye Nile Red (Sigma #72485) was dissolved in peanut oil (final concentration 20 µg/ml). Fast green (Sigma #F7258) to stain proteins (first dissolved in distilled water at 1mg/ml) was added to the diluted milk or syrup (final concentration 60 µg/ml). SOE and MOE emulsions were prepared using solutions as described above and samples of each were diluted and placed on a glass slide, covered with a coverslip, and imaged immediately using an inverted confocal microscope (Leica DMI6000 AFC, Model SP8)^83^. Fluorescence microscopy images of the emulsions were processed using ImageJ v1.53t^84^.

### Administering formulations to mice

To reduce handling stress^85^, transparent training tunnels (Zoonlab #3084094) were used as described^34^ for transferring and identifying the mice (Fig. S4). Training and treatment of mice were as published^7^, with the following modifications. The formulations were offered to the mice using a variable volume type P200 PIPETMAN® P (Gilson) with a sterile 200 µl graduated filtered plastic tip (TipOne® # S1120-8810, Starlab, Germany). Mice were collected from the home cage and transferred to the grid of a second cage, and gently held by the tail. The formulations were offered by dropwise expelling the liquid from the tip of the pipette as soon as the mouse started to drink. Consumption time was recorded from the moment the liquid was offered until all the liquid from the tip was consumed. If, after 60 seconds, the mouse had not drunk or had drunk only part of the formulation, it was restrained by gently grabbing the scruff of its neck and the remaining formulation was offered again. Total consumption time is the time before restraining plus, if applicable, the time to consume the remaining formulation with restraining. Treated mice were temporarily placed in a 3^rd^ cage and then returned to the home cage when all the mice of a cage were treated. For the treatment with TAM containing emulsions, mice received a dispensed volume that was adjusted based on the weight of the mice as recorded during the last training day. For OG, a weight-adjusted OIL-TAM solution was prepared for each mouse by diluting the same 100 mg/ml TAM stock solution used to make the MOE and SOE emulsions. The corresponding mouse then received 200 µl using a curved OG needle of 50 mm/18 G (Finescience, #18061-50) attached to a 1 ml syringe. Non-treated control mice or mice treated with different formulations or dosages were kept in separate cages to prevent confounding effects due to coprophagy.

### Flow cytometer analysis of ex vivo cells

Mice were euthanized 40 h after the last TAM treatment and single-cell suspensions were prepared from each thymus. For HY-switch mice, cells were stained with AQUA Zombie live/dead dye (Biolegend #423101) according to the manufacturer’s instruction and then surface stained for TCR β (BioLegend #2629564), CD8a (BioLegend #2562610), CD4 (BioLegend #893330), TCR H-Y (eBioScience #466267), CD3e (eBioScience #469315), NK1.1 (BioLegend #313312), CD19 (BioLegend #313641), CD25 (BioLegend #313392) in the presence of an Fc receptor blocking antibody (BioLegend AB_1574973) during 30 min at 4°C. For tdTomato expression, cells were stained with Zombie NIR live/dead dye (Biolegend #423105) and then surface stained for TCR β, CD3, CD8, and CD25 as above, in addition to CD4 (Biolegend #100428). After washing, samples were acquired with an LSR II Fortessa cytometer (BD Biosciences, USA) and analyzed by manual gating using FlowJo™ v10.4 Software (BD Life Sciences). For HY-switch cells, gates were set to exclude dead cells, doublets, and lineage (NK1.1, CD19, and CD25) positive cells. HY-TCR expression was analyzed by setting a gate based on cells from non-treated mice. For R26-CreERT2-Ai14 mice, a gate was set to exclude dead cells and doublets. tdTomato expressing cells were quantified using a gate based on non-treated cells.

### Serum collection

Mice were euthanized and blood samples collected via cardiac puncture using 25 G needles (Terumo #AN*2516R1). Blood was transferred to SST tubes (BD Microtrainer® #365968) and processed according to the manufacturer’s instructions. Individual sera were stored frozen until analysis.

### Tissue collection and microscopic analysis

Spleen tissues were fixed in 4% formaldehyde overnight at 4°C and then cut with a microtome (Micron KS34 ThermoFisher) to obtain 30 μm tissue sections^86^ that were then stained for 10 min with 0.3 µM DAPI in DPBS (Thermo Scientific™ #62247). The sections were mounted with Fluoromount-G™ mounting medium (Invitrogen #00-4958-02) and analyzed with a slide scanning microscope (Vectra Polaris PerkinElmer, USA). Images were acquired and processed using ImageJ and QuPath v0.3.0^87^ using identical settings for each channel.

### Statistics and data analysis

The non-inferiority test was performed based on published guidelines^38^. The 95% confidence interval of the difference between means (CI) was derived using Dunnett’s test to correct for the multiple comparisons of multiple datasets to a single control dataset^88, 89^.

As non-inferiority margin, we choose 1.5 times the standard deviation (SD) of the control group. One-way ANOVA followed by Dunnett’s test was applied when comparing multiple datasets to a single control dataset. All statistical analyses were computed using Prism (v9.2.0 GraphPad) and, where applicable, a p-value < 0.05 was considered statistically significant.

Tukey box plots in graphs show the minimum and maximum values (ends of the whiskers), interquartile range (length of the box), and median (line through the box) of sets of data. Individual data points are shown as circles.

Power calculations to determine the group size for the non-inferiority test were performed in the statistical environment R, using a one-tailed t-test with a power of 0.8 and a significance level (alpha) of 0.025.

## Acknowledgements

We are grateful for helpful discussions and practical support by Dr. Sina Schalbetter, Dr. Philipp Schätzle, Hafida Atiqi and the animal facility of the UZH. We acknowledge Dr. Tina Fabia Notter for her valuable input and suggestions in reviewing and editing our manuscript. We appreciate Zoolab for donation of samples of training tunnels. We thank the late Prof. Michal Denison and Dr Guochun He of the University of California, Davis (USA) for providing the VM7Luc4E2 cell line as well as the relevant protocols to work with these cells.

## Data availability

The data underlying examples and figures are available from the corresponding author upon reasonable request.

## Author contribution

DV, VB, and TB wrote the manuscript. DV and TB conceptualized and supervised the work. VB and DV performed the experiments.

---

**Figure S1:**
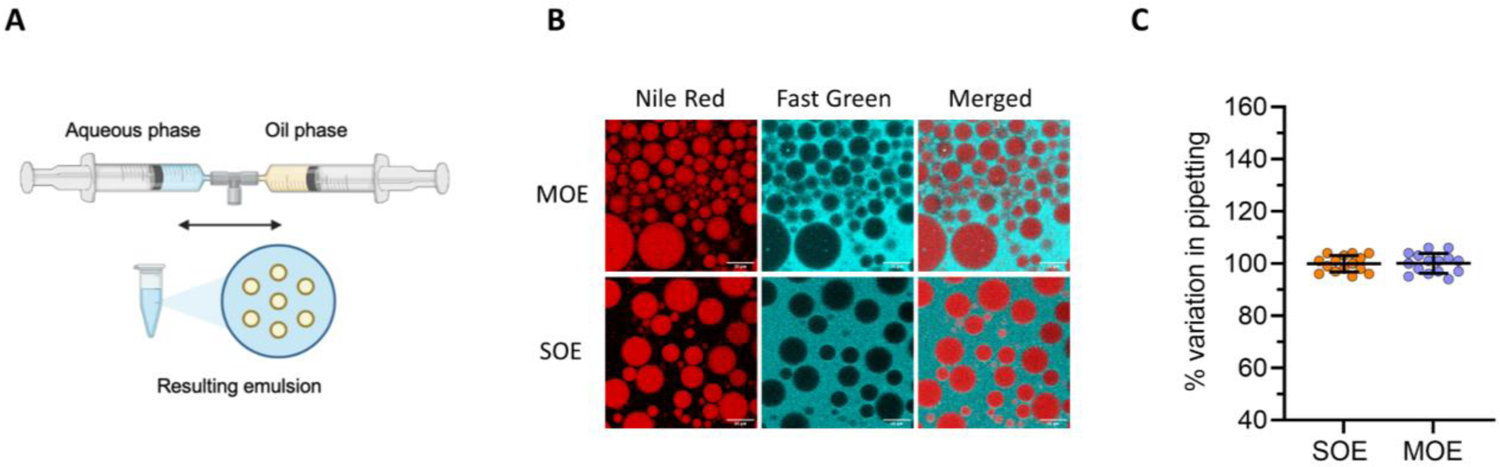
Oil-based emulsions as vehicles for TAM administration. (A) The 2-syringe homogenization method involves connecting two syringes via a suitable adapter. The two solutions (aqueous and oil) are then mixed by repetitively pushing the plungers until a homogeneous emulsion with dispersed oil droplets is obtained. Image created with Biorender. (B) Representative fluorescence microscopy pictures of MOE and SOE. Oil drops are stained with Nile Red (red) and proteins in the milk or syrup are stained with Fast Green (Cyan). Magnification 20x. White scale bar=20 µm. (C) 75 µl of MOE (blue) or SOE (orange) emulsions were pipetted using the reverse pipetting approach and weighed (n=20). Depicted is the % of variation in pipetting (relative to the average weight) ±SD for each formulation.

**Figure S2:**
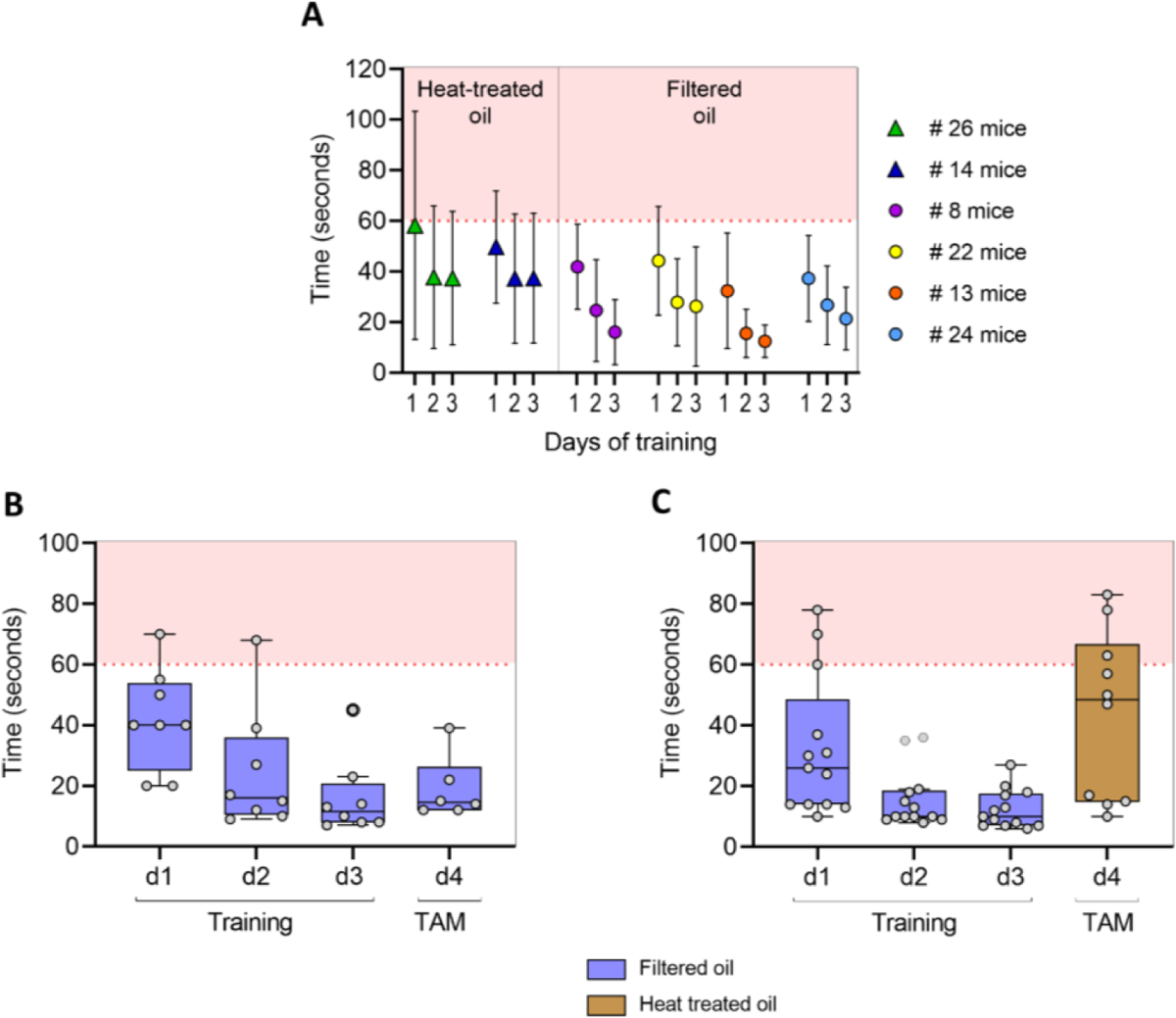
Palpability of emulsions made with filtered or heat-treated oil. (A) Compilation of consumption times for three days of training, recorded and depicted as described in Figure 1E. Data is from different experiments where mice were offered sweetened milk-oil emulsions (MOE) made with either heat-treated oil (triangles) or sterile filtered oil (circles). Each experiment is indicated with a different color and the number of mice per experiment is indicated in the legend. The mean consumption time ± SD is indicated for each training day. (B and C) Consumption times during training with MOE made with filtered oil (blue bars) and MOE-TAM treatment made with emulsions containing filtered oil (B) or heat-treated oil (C).

**Figure S3:**
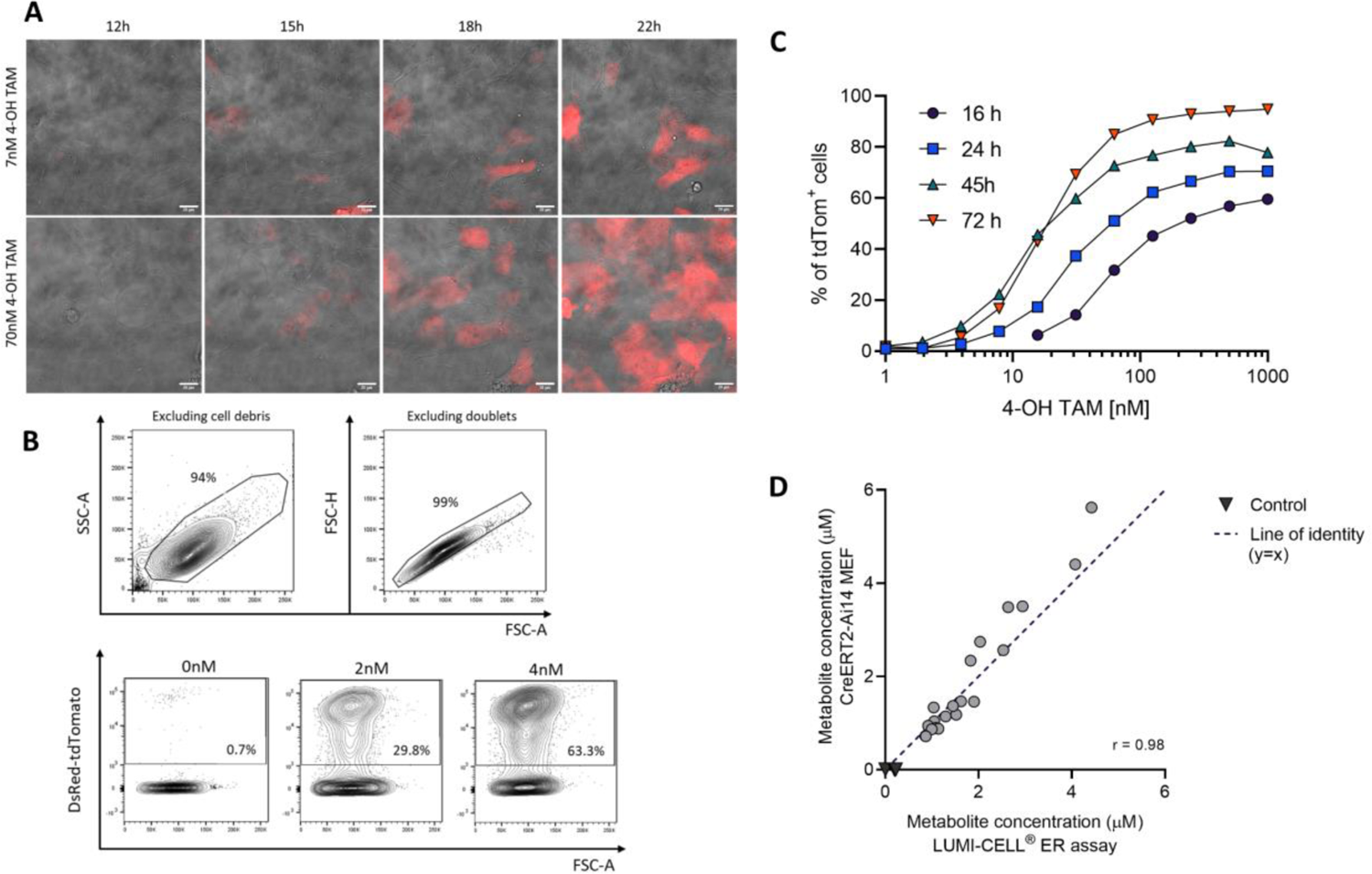
Specifics of an in vitro assay for the quantitative detection of serum TAM metabolites. (A) R26-Cre-ER^T2^ Ai14 mesenchymal embryonic fibroblast cells (MEFs) were cultured and treated with 7 nM (top panels) or 70 nM (bottom panels) 4-OH TAM for 28 hours. Induction of tdTomato expression (red) was followed over time by taking pictures (bright field and fluorescence) at the indicated time points with magnification 63x. Scale bar = 50 µm. (B) Representative dot plots of a flow cytometric analysis of R26-Cre-ERT2 Ai14 MEFs treated with 0 nM, 2 nM or 4 nM OH-TAM for 45 hours. Gates on the dot plots were set to exclude cell debris and doublets (top panels) and to calculate the percentage tdTomato expressing MEFs (bottom panels). (C) Percentage of tdTomato expressing cells, analyzed as shown in (B), following stimulation with a range of 4-OH TAM concentrations (1-1000 nM, 2-fold dilution) for the indicated time periods (1-72 h). Each data point is the average of independent duplicate measurements. (D) TAM-metabolite concentrations in sera from mice collected 6 h after TAM treatment, were determined using R26-Cre-ERT2 Ai14 MEFs and compared to that obtained using an independent LUMI-CELL ER assay. MEF cells were cultured in the presence of the different mouse sera for 45 hours and the percentage of tdTomato positive cells quantified as shown in (B). The concentration was calculated based the response of the cells to a range of 4-OH TAM concentrations, resulting in a standard curve similar as shown in (C). Results obtained with both R26-Cre-ER^T2^ Ai14 MEFs and the LUMI-CELL ER assay were compared for each serum sample. A line of identity (y=x) was inserted as a reference for similarity and the correlation coefficient r was calculated.

**Figure S4:**
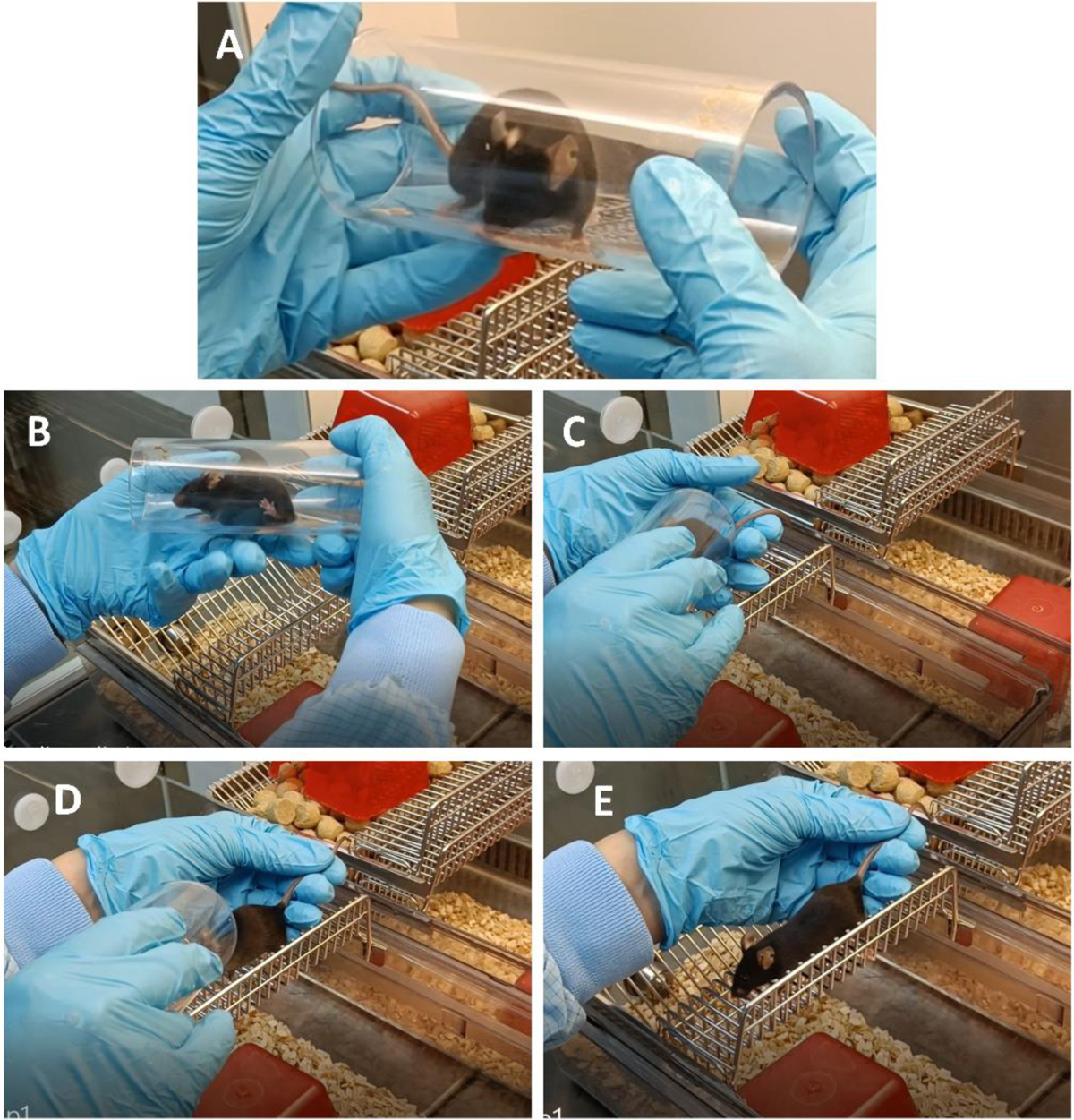
Handling of mice using training tunnels. Mice were habituated to tunnels for at least two days prior to training and treatments. (A) Mice were collected from the home cage using the tunnel and identified based on earmarks. (B-E) The animals were transferred to the grid of an empty cage and offered a formulation with a micropipette while gently held by the tail (Fig 1B).

## Supplementary Video 5A and 5B

**Videos demonstrating voluntary and restrained consumption of milk-oil-emulsions**. Experimental setup for training and TAM treatment of mice includes 2 new cages (left) and the home cage (right). Mice are collected from the home cage and placed on the new cage using a tunnel as shown in Fig S4. <u>Video 5A:</u> Representative example of voluntary consumption. The mouse is gently held by the tail with one hand and offered the solution using a 200 µl pipette with the other hand. Once the mouse shows interest in the offered solution, small volumes are expelled according to how fast the mouse drinks the solution. <u>Video 5B:</u> Representative example of restrained consumption. The mouse is gently restrained by holding the scruff of the neck with one hand while it is offered the formulation with the other. Also here, small volumes are expelled from the pipette tip according to how fast the mouse drinks the solution.

